# Matching lynx population estimates to management scale: conservation science vs politics in the western Swiss Alps

**DOI:** 10.64898/2026.08.25.747176

**Authors:** Stijn Verschueren, Veronika Braunisch, Valentin Debons, Raphaël Arlettaz

## Abstract

Reliable population estimates are essential for wildlife management, yet monitoring schemes often do not match the administrative scale at which decisions are made. We illustrate this challenge using the Eurasian lynx in the canton of Valais, Switzerland, where official state monitoring is fragmented across three reference areas surveyed in different years. We analyzed three winters (2023–2026) of independent, canton-wide camera trap data, recording 899 independent lynx captures (27, 31 and 34 adults per winter). Lynx distribution and reproduction concentrated in the Northwest and connected to the thriving Pre-alpine populations. Density modelling for 2025/2026 estimated 37 independent lynx (95% CI: 26–52) on the whole cantonal territory, corresponding to a density of 1.09 (0.77–1.56) individuals per 100 km^2^. These estimates are substantially below the figures improperly extrapolated from a cross-cantonal reference area and conveyed by political authorities. Future lynx management decisions should be rooted in scientifically sound, scale-relevant information.

## INTRODUCTION

The Eurasian lynx (*Lynx lynx*) was originally a wide-ranging apex predator of Eurasian forest ecosystems, but was extirpated from much of western Europe during the nineteenth and twentieth centuries (Breitenmoser & Breitenmoser-Würsten, 2024). Reintroduction programs since the 1970s have re-established populations in several regions, but many remain small and isolated (Linnell et al., 2009). Population expansions are constrained by multiple interactive factors, including habitat fragmentation, infrastructure-induced mortality, illegal hunting and the genetic constraints typical of small founder populations (Borel et al., 2026; Vogt et al., 2025). Their individual effects are difficult to disentangle, which represents an impediment to proper management. To all intents and purposes, effective wildlife conservation depends on evidence-based management informed by reliable monitoring schemes yielding correct demographic information (Nichols & Williams, 2006).

Switzerland has played an important role in the restoration of the Eurasian lynx in western Europe and subsequently developed extensive expertise in its monitoring and population ecology (KORA, 2022). The country established a systematic national monitoring program based primarily on standardized camera trap surveys. Twelve reference areas are surveyed periodically, with individually identifiable pelage patterns providing the basis for spatially explicit capture–recapture estimates of density and abundance. This led to the most recent national estimate of 364 (± 10) independent lynx in 2024/2025 (https://www.kora.ch/en/species/lynx/abundance). Despite this encouraging recovery, lynx remains a national priority species in Switzerland as major conservation threats persist (BAFU, 2025).

The Swiss reference areas have been designed around established and emerging lynx (sub)populations. This is ecologically appropriate for monitoring population status and trends, but these survey units do not match the administrative scale (the cantons) at which management decisions are made. The canton of Valais illustrates the consequences. Although lynx occur in Valais since the late 1970s, following reintroduction efforts initiated in Central Switzerland in 1971 (Breitenmoser & Breitenmoser-Würsten, 2008), only 15 adult lynx (with a maximum of 8 per winter) occupied its northern part between 2011 and 2015, at an estimated density 0.32 individuals per 100 km^2^ (Biollaz et al., 2015). Moreover, the species had in between mysteriously disappeared from most of the Pennines Alps, south of the Rhône, where lynx were still roaming numerous in 1985-1988 (Haller, 1992). Long suspected, illegal persecution was first officially documented in 2015 when a dense network of ejecting-strangling snare traps was discovered along an important dispersal corridor linking Valais with the Swiss Pre-alps where lynx thrive (Arlettaz et al., 2021). With the conviction of a hunter and partial relief of this poaching pressure due to a wide media coverage, lynx occurrence increased in parts of the canton, especially in the Northwest, but recovery remained spatially heterogeneous and extensive suitable habitat is still left unoccupied, while illegal shooting with hunt guns is still occurring (Arlettaz, 2026b, 2026a).

This heterogeneity is reflected in recent density estimates from the three state monitoring reference areas that overlap Valais (Fig. 1). These areas are surveyed as part of the official monitoring program, conducted by KORA (Swiss Coordination for Carnivore Ecology and Wildlife Management) and commissioned by the Federal Office for the Environment in collaboration with local state game wardens. State monitoring detected no lynx in reference area IVd during winter 2018/2019; density in reference area IVe was estimated at 1.25 independent lynx per 100 km^2^ (95% CI: 0.99–1.51) in winter 2024/2025; and density in reference area IVc reached 5.12 individuals per 100 km^2^ (4.04–6.19) in winter 2021/2022 (https://www.kora.ch/en/species/lynx/camera-trap-monitoring). Noteworthy, the latter area extends much beyond Valais into the Swiss Pre-alps (cantons of Vaud and Bern), where lynx densities are among the highest in the country. Because the official monitoring estimates refer to different areas surveyed in different years, they cannot be directly combined or extrapolated to derive a robust canton-wide population estimate for Valais. Nevertheless, such interpretations have appeared in public and policy discussions, initially conveyed by cantonal political authorities, with claims that Valais as a whole may support as many as 80 lynx, and at a density of more than 5 lynx/100 km^2^ north of the Rhône river (Chillier, 2026; DJFW Canton of Valais, 2025; Zengaffinen, 2025).

**Fig 1.**
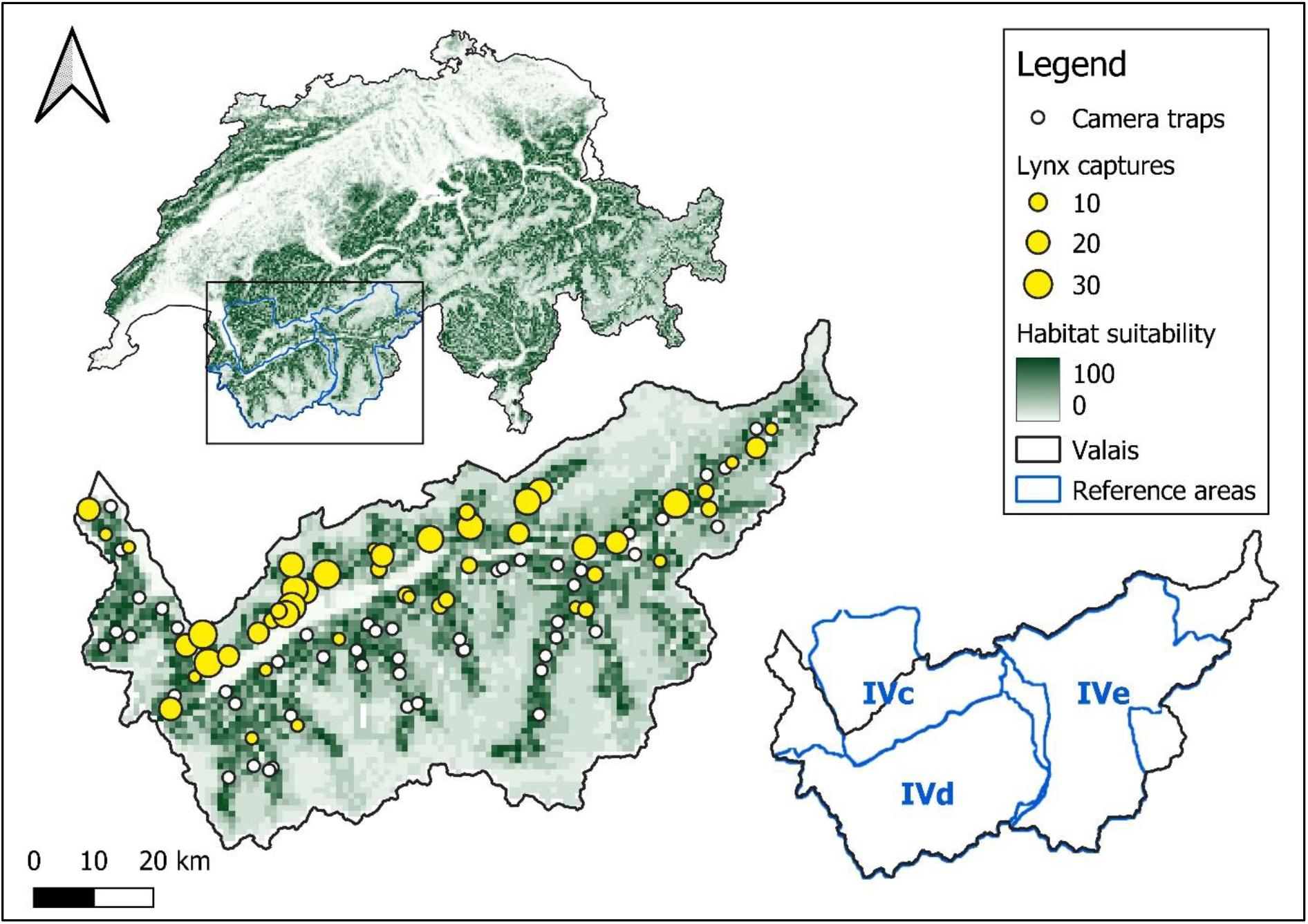
Map of the lynx captures and camera stations deployed by the Conservation Biology Division of the University of Bern in the canton of Valais, Switzerland during the winter 2025/2026 with the number of lynx captures (a white dot indicates no lynx capture) and habitat suitability (derived from Zimmermann 2004) shown in green shading. The reference areas of the official state monitoring are outlined in blue: IVc = North of the Rhône (note that ca 65% of lynx habitat lies outside of Valais, in the cantons of Vaud and Bern); IVd = Southern Lower Valais; IVe = Upper Valais. Each of these reference areas is in principle monitored every third year.

The need for management-relevant population estimates has become increasingly urgent as Switzerland debates the regulation of large carnivore populations, first for wolves and now for lynx (Papaux, 2026b, 2026a). Decisions regarding population regulation require reliable information at the administrative scale where management actions are implemented. In the absence of such information, extrapolations from ecologically defined monitoring units may convey a misleading impression of local abundance and weaken the evidential basis for appropriate regulatory decisions.

Long-term camera trap monitoring by the Conservation Biology Division of the University of Bern offers an independent opportunity to address this information gap. Since 2011, 102 cameras have been deployed annually across the canton during winter in order to investigate the spatio-demographic interactions between recovering large carnivores and their ungulate prey. Here, we report the results of the three most recent winter surveys (2023/2024, 2024/2025, 2025/2026) in terms of lynx presence, distribution, reproduction and minimum number of individuals identified, and apply a spatially explicit capture–recapture (SECR) model to the 2025/2026 data to estimate density and abundance across Valais and within the three official reference areas. While analysis of the full time series is ongoing, this interim assessment provides urgently needed population information at the scale at which regulation is currently being discussed. It further highlights a broader challenge: delivering scientific evidence in time to meet emerging policy discourses.

## METHODS

### Study area and camera trap monitoring

The canton of Valais lies in the western Swiss Alps, structured by the deep Rhône valley: a densely populated valley floor flanked by forested mid-elevation slopes (500–2,300 m a.s.l.) that grade into alpine meadows, rocky outcrops and permanent snow and glaciers reaching 4,634 m a.s.l.. Forest cover is mainly coniferous and mixed stands interspersed with grassland, settlements and roads. Topography, human infrastructure and winter snow accumulation constrain wildlife movement and concentrate activity within the mid-elevation forested belt, particularly in winter when high-elevation habitats are largely unused (Fig. 1).

We conducted independent standardized camera trap surveys each winter since 2011 and use data from the three most recent winters (2023/2024–2025/2026) for the present assessment. Camera stations were distributed on a 10 × 10 km grid over approximately 1,900 km^2^ forested study area, with three sites targeted per cell and stations spaced 2–3 km apart along forest trails. Station locations were held constant across winters. Over the three assessment winters we operated a mean of 99 stations per winter (range 98–101) from the network of 102 permanent stations, the annual total varying only through theft or vandalism. Each station comprised one motion-activated camera (Reconyx PC900 HyperFire Professional IR or HP2X Professional; Reconyx, Wisconsin, USA) mounted 40–60 cm above ground along forest trails or unpaved roads, capturing three-image bursts per trigger (1 s refractory interval) and operating for a mean of 124 days per winter.

### Individual identification

Images were classified to species in Lepus Pro (https://lepus.pro). Lynx were identified individually from their unique pelage patterns. Because cameras were single-sided, we first compiled separate right- and left-flank catalogues, then matched flanks across sides using spatial and temporal co-occurrences of paired detections, together with coat-pattern type, sex and body size. Matches were assigned only when all criteria were met; otherwise records were retained as single-flank identifications. Assignments were supported using ArgusWild AI (https://www.arguswild.ai), a computer-assisted pattern-recognition platform requiring human confirmation of every match (Verschueren et al., 2023). Detections that could not be confidently assigned to an individual were retained as unidentified lynx records and we report identification success rate per winter.

### Analysis

For each of the three winters we report the number of independent lynx captures (i.e., independent if they involved different individuals, or the same individual photographed more than 30 min apart), the naïve occupancy (i.e., the proportion of trail camera stations recording lynx), the number of adult individuals identified (i.e., both flanks, and left- and right-flank only), the number of litters and cubs, and the identification success rate (i.e., the proportion of captures confidently assigned to an individual).

We estimated lynx density and abundance for the 2025/2026 survey using a spatially explicit capture– recapture model (SECR), with a half-normal detection function (Efford, 2024). Detection histories were constructed over a three-month period (December – February) with daily sampling occasions running from midday to midday. We considered all individuals that had both sides matched and individuals with right sides only. We excluded three individuals that were detected outside the SECR time window.

We fitted a canton-wide model assuming spatially constant density, with the habitat mask restricted to areas with a habitat suitability index ≥0.2 (on a scale of 0–1) within the cantonal boundary (Zimmermann, 2004). We further summarize the lynx detections and survey effort for the 2025/2026 winter within the three official reference areas (full coverage of IVd an IVe, 35% coverage of IVc), and applied the same density model to three subsets of camera sites grouped by reference area. In addition, we calculated an apparent density for each reference area as the number of individually identified lynx divided by the area of suitable habitat. This uncorrected metric illustrates how densities may be overestimated when the focal area is small and when the political boundaries or the boundaries of the reference monitoring areas do not constrain lynx movements, in such a way that individuals detected near the boundaries may also range outside the focal area.

## RESULTS

### Detections and individual identification

Over the three winters we recorded 899 independent lynx captures. Captures were comparable in the first two winters but declined in 2025/2026, despite the highest survey effort of the period (12,735 camera-nights). Naïve occupancy rose from 0.39 in 2023/2024 to 0.49 in each of the two subsequent winters, and the minimum number of identified adult individuals increased steadily across the three winters, from 27 to 31 to 34. Reproduction was documented in every winter, with 8 litters (at least 11 cubs) in 2023/2024, 7 litters (at least 11 cubs) in 2024/2025 and 4 litters (at least 7 cubs) in 2025/2026. Identification success was consistently high, with at least 87% of lynx captures assigned to an individual in every winter (Table 1).

**Table 1.** Summary of trail camera survey effort and lynx detections in the canton of Valais across three winters (2023/2024–2025/2026).

| Winter | 2023/2024 | 2024/2025 | 2025/2026 |
| --- | --- | --- | --- |
| Camera stations | 101 | 99 | 98 |
| Camera nights | 11,787 | 12,476 | 12,735 |
| Stations with lynx | 39 | 49 | 48 |
| Naïve occupancy | 0.39 | 0.49 | 0.49 |
| Independent lynx captures | 319 | 324 | 256 |
| Minimum number of adult individuals | 27 | 31 | 34 |
| Adults with both flanks | 21 | 22 | 25 |
| Adults with left flank only | 6 | 9 | 8 |
| Adults with right flank only | 3 | 4 | 9 |
| Litters | 8 | 7 | 4 |
| Cubs (minimum) | 11 | 11 | 7 |
| Identification success rate | 87% | 89% | 90% |

### Density and abundance (2025/2026)

The canton-wide model estimated an overall density of 1.09 (95% CI: 0.77–1.56) independent lynx per 100 km^2^. The detection probability was 0.018 (95% CI: 0.014–0.023), and the movement parameter was 4.07 km (95% CI: 3.64–4.54). Applied across the habitat mask, the abundance estimated by the model was 37 independent lynx (95% CI: 26–52) within Valais during the winter 2025/2026.

Lynx were concentrated in reference area IVc, north of the Rhône, which is connected with the Pre-alpine population. In 2025/2026 in IVc, we captured 139 out of 256 lynx captures and 18 out of 34 individuals, as well as all litters (n = 4) and cubs (n = 7). Reference areas IVd and IVe were only partially occupied, with 6 and 10 individuals identified, respectively, and no evidence of reproduction. Based on the number of individuals identified over the area of suitable habitat, the apparent densities were 6 (IVc), 0.58 (IVd) and 0.88 (IVe) individuals per 100 km^2^. However, accounting for imperfect detection and allowing activity centers to occur beyond reference area boundaries resulted in a substantially lower density estimate for IVc, with 1.74 individuals per 100 km^2^ (95% CI: 1.03–2.93). Modeled density was slightly higher in IVe 0.93 (0.49–1.76), while in IVd, the number of captures and recaptures was too low to obtain a reliable density estimate and associated confidence intervals (Table 2).

**Table 2.**
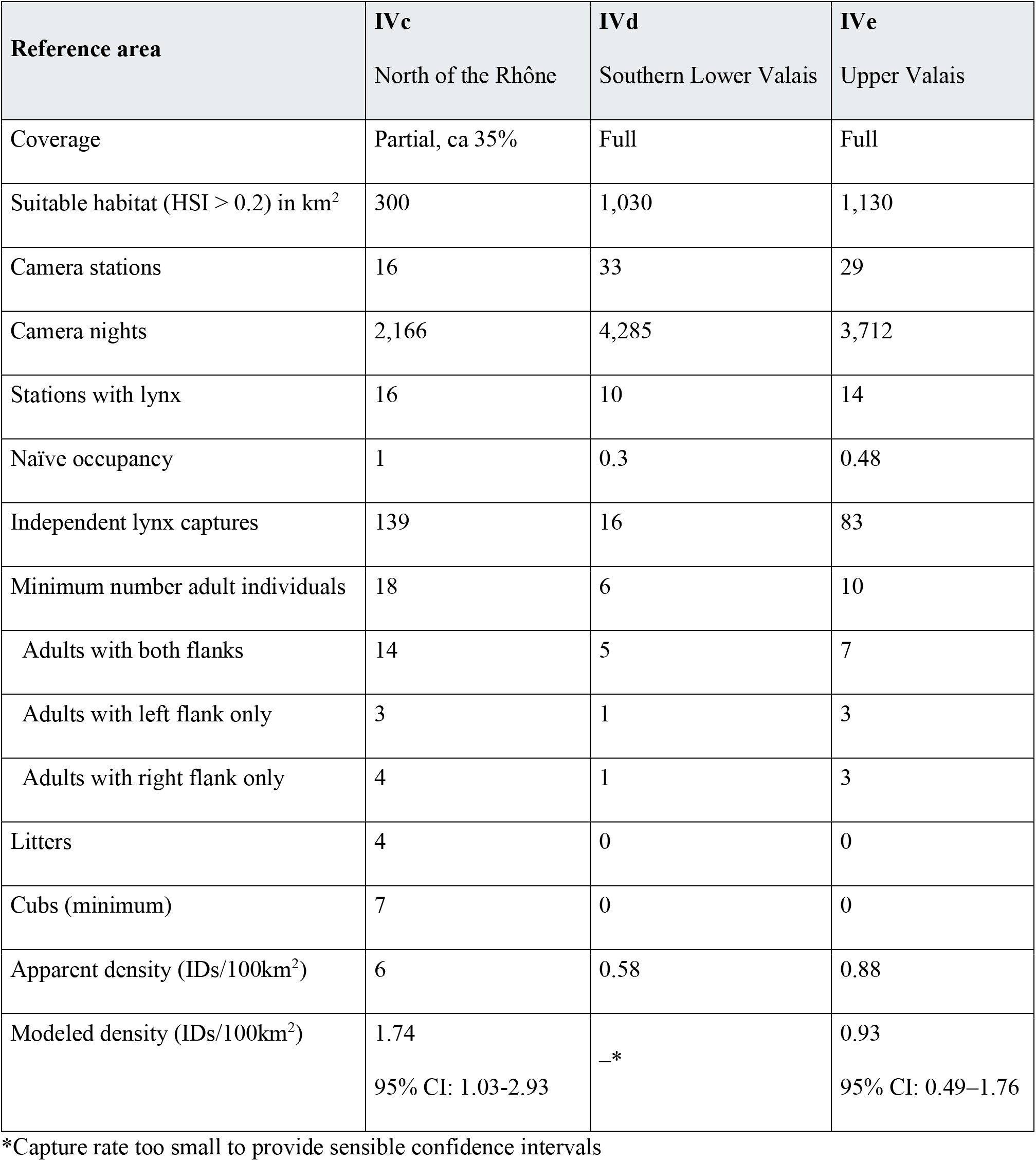
Summary of survey effort and lynx detections during winter 2025/26 in the three reference areas that overlap the canton of Valais. Coverage indicates whether our camera trap array spanned the full reference area.

## DISCUSSION

We estimated 37 independent lynx (95% CI: 26–52) over 3,400 km^2^ suitable habitat in Valais during the winter 2025/2026, at an overall density of 1.09 (0.77–1.56) individuals per 100 km^2^. The modeled density estimate for the Valais part of reference area IVc, where lynx concentrated, was substantially lower (ca. one third) than the 2021/2022 value reported for the full reference area (5.1), despite the population size having increased in Valais in the intervening period. Our abundance estimate, and its upper confidence limit of 52 individuals, is also well below the claim of up to 80 individuals circulating in media outlets. Thus, the figures put forward by the political authorities, notably in anticipation of a possible future regulation of lynx in the area IVc, appear to rely on extrapolations from data that do not originate exclusively from Valais. Whether this overestimation of abundance and density reflects a mis-interpretation of the evidence at hand or a deliberate strategy remains unclear. This notwithstanding, management decisions by the cantonal and federal states must be taken on monitoring data reflecting the real situation of lynx in Valais as their appropriate evidential basis.

The apparent density of 6 individuals per 100 km^2^ in reference area IVc further illustrates the potential for substantial overestimation when identified individuals are scaled to a relatively small area of suitable habitat (∼300 km^2^, Table 2) without accounting for movements beyond its boundaries. Lynx are wide-ranging, and individuals detected in the northern Valais part of IVc move beyond this area, as evidenced by repeated recaptures outside this area, and with model derived densities being three times lower.

Apparent and modelled densities were more similar in reference area IVe, where the modelled estimate (0.93) was slightly higher than the apparent density (0.88). The fewer number of individuals, the larger spatial extent of this area (∼1,130 km^2^) and topographical constraints to movement beyond the boundaries reduces the influence of edge effects, so that apparent density under these conditions present a better approximation of the true density. This is further supported by the fact that our density estimate was close to, albeit slightly below, the value reported by KORA for 2024/2025 (1.25). The number of captures in reference area IVe was too low for a reliable SECR estimate, yet the detection of lynx in this area indicates an increase and range expansion relative to the KORA survey in 2018/2019, when no lynx were detected. This is corroborated by our own findings since 2011, and is an encouraging sign for the conservation and restoration of a nationally threatened carnivore.

Notably, our modelled density estimates for the two reference areas had wide confidence intervals, reflecting the relatively limited number of cameras and individual recaptures within each reference area. Our survey was designed to provide representative coverage across the entire canton rather than within each reference area. Our canton-wide estimates anyways provides the most robust basis currently available for assessing the current lynx population in Valais.

Suitable habitat remains widespread across Valais, yet our results show that lynx remain spatially unevenly distributed, with persistent low occurrence in poorly connected parts of the canton such as the southern valleys (Pennines Alps), from where they have been (illegally) eradicated in the 1990s and 2000s (Huysecom, 2025). Exploratory analyses indicate that lynx density appears highly structured by the ecological distance from the dense population in the Pre-alps, encompassing the reference area IVc. This patterns suggests that connectivity, rather than habitat availability, constrains lynx dispersal and therefore expansion, and this pattern is reinforced by the history of illegal persecution along the corridor linking Valais to the Pre-alps (Arlettaz et al., 2021). A forthcoming analysis of the full time series from 2011 will incorporate connectivity and habitat suitability to model this structure explicitly and to assess recovery over a 15-year period.

The expansion of the Valais lynx population since Biollaz et al. (2015) points to a spatially concentrated phase of recovery in recent years. Populations released from poaching pressure and occupying vacant habitat typically grow steeply before logistic density dependence regulation takes hold, and the increase is concentrated where colonizers arrive first (Pletscher et al., 1997). Such localized increases are therefore expected and do not indicate that the canton as a whole is nearing carrying capacity: large areas of suitable habitat remain poorly colonized if not void of lynx, with over half of camera stations not recording any lynx. A rapid local increase and high apparent density can nonetheless create the perception of a population growing beyond acceptable levels. Distinguishing a canton-wide trend from locally concentrated recovery is therefore central, and we recommend to anticipate the population’s future trajectory rather than curtailing an expansion that remains far from its historic distribution. Doing so requires evidence matched to decisions in both scale and timing. The difficulty is that regulation is debated during early expansion, when provisional figures spread faster than analyses can correct them. For now, we suggest leaving the Valais lynx population freely recover, further expanding into the area south or the Rhône where it was still roaming in numbers some decades ago. We also advise to envision regulation by culling operations only once overall carrying capacity is achieved throughout Valais. This way Valais would eventually play its key node role for reinstating a robust international, Alpine metapopulation system with functional connections to Northern Italy and Eastern France, where the species is still scarce due to poor dispersal from the Northern Alps.

## REFERENCES

Arlettaz, R. (2026a). Étrange fin pour un lynx et mise en scène macabre / Rätselhaftes Ende eines Luchses und Hinweise auf makabre Inszenierung. Fauna.vs Info, 10–15.

Arlettaz, R. (2026b). Le lynx en Valais: la régulation officielle supplantera-t-elle la longue tradition de régulation illégale? / Der Luchs im Wallis: Wird die offizielle Bestandsregulierung die lange Tradition der illegalen Regulierung ablösen? Fauna.vs Info, 26–31.

Arlettaz, R., Chapron, G., Kéry, M., Klaus, E., Mettaz, S., Roder, S., Vignali, S., Zimmermann, F., & Braunisch, V. (2021). Poaching threatens the establishment of a lynx population, highlighting the need for a centralized judiciary approach. Frontiers in Conservation Science, 2. 10.3389/fcosc.2021.665000

BAFU. (2025). Digitale Liste der National Prioritären Arten.

Biollaz, F., Mettaz, S., Zimmermann, F., Braunisch, V., & Arlettaz, R. (2015). Status du lynx en Valais quatre décennies après son retour: suivi au moyen de pièges photographiques. Bulletin de La Murithienne, 133, 29–44.

Borel, S., Marti, I., Origgi, F. C., Delalay, G., Breitenmoser, C., Zürcher-Giovannini, S., Frey, C. F., Basso, W., Schweizer, D., Kittl, S., Ryser-Degiorgis, M.-P., & Keller, S. (2026). Causes of morbidity and mortality in free-ranging Eurasian lynx (Lynx lynx) in Switzerland, 2000–2022. PLOS One, 21(3), e0344107. 10.1371/journal.pone.0344107

Breitenmoser, U., & Breitenmoser-Würsten, C. (2024). Eurasian Lynx Lynx lynx (Linnaeus, 1758). In K. Hackländer & F. E. Zachos (Eds.), Handbook of the Mammals of Europe. Springer. 10.1007/978-3-319-65038-8_123-1

Breitenmoser, U., & Breitenmoser-Würsten, C. (2008). Der Luchs - Ein Grossraubtier in der Kulturlandschaft. Salm Verlag.

Chillier, G. (2026). Le Valais tout près de réguler le lynx. Le Nouvelliste. 12 June 2026.

DJFW Canton of Valais (2025). Infoblatt Luchs 2025. https://www.vs.ch/de/web/scpf/fiche-d-informations-especes

Efford, M. (2024). secr: Spatially explicit capture-recapture models. R Package Version 4.6.5.

Haller, H. (1992). Zur Ökologie des Luchses Lynx lynx im Verlauf seiner Wiederansiedlung in den Walliser Alpen. Mammalia Depicta, 15(62).

Huysecom, L. (2025). Construction et déconstruction d’un conflit entre humains et prédateurs : le cas de la réintroduction du lynx en Val d’Anniviers. Master of Science in geography, University of Lausanne. 137 pages.

KORA. (2022). 50 years of lynx presence in Switzerland.

Linnell, J. D. C., Breitenmoser, U., Breitenmoser-Würsten, C., Odden, J., & von Arx, M. (2009). Recovery of Eurasian Lynx in Europe: What Part has Reintroduction Played? In Reintroduction of Top-Order Predators (pp. 72–91). Wiley. 10.1002/9781444312034.ch4

Nichols, J., & Williams, B. (2006). Monitoring for conservation. Trends in Ecology & Evolution, 21(12), 668–673. 10.1016/j.tree.2006.08.007

Papaux, S. (2026a). «Des voix s’élèvent en Valais» pour tirer les lynx. Watson. 4 August 2026.

Papaux, S. (2026b). Le retour du lynx en Valais révèle des zones d’ombre. Watson. 10 June 2026.

Pletscher, D. H., Fritts, R. H., & Ream, R. R. (1997). Population Dynamics of a Recolonizing Wolf Population. The Journal of Wildlife Management, 61(2), 459–465. 10.2307/3802604

Verschueren, S., Fabiano, E. C., Kakove, M., Cristescu, B., & Marker, L. (2023). Reducing identification errors of African carnivores from photographs through computer-assisted workflow. Mammal Research, 68, 121–125. 10.1007/S13364-022-00657-Z

Vogt, K., Korner-Nievergelt, F., Signer, S., Zimmermann, F., Marti, I., Ryser, A., Molinari-Jobin, A., Breitenmoser, U., & Breitenmoser-Würsten, C. (2025). Long-term changes in survival of Eurasian lynx in three reintroduced populations in Switzerland. Ecology and Evolution, 15(4). 10.1002/ece3.71095

Zengaffinen, N. (2025). Der Luchs erobert das Oberwallis: Fünfmal mehr Exemplare als vor 5 Jahren. Walliser Bote. 15 June 2026.

Zimmermann, F. (2004). Conservation of the Eurasian Lynx (Lynx lynx) in a fragmented landscape – habitat models, dispersal and potential distribution. University of Lausanne.

